# Compound signaling activates endogenous retroviruses by inducing enhancer and gene-neighborhood transcription

**DOI:** 10.1101/284695

**Authors:** Saliha Azébi, Eric Batsché, Fréderique Michel, Etienne Kornobis, Christian Muchardt

## Abstract

**Summary:** Multiple sclerosis (MS) is a neuroinflammatory and autoimmune disease, in which various immune cell types and autoreactive T cells exert a pathogenic activity. This disease is also associated with increased transcription of several endogenous retroviruses (HERVs) normally kept in check by heterochromatin. Here, we have uncovered an organic pollutant dieldrin that activates several HERVs associated with MS and allowing us to examine the mechanism of their activation. Dieldrin singles out by its ability to simultaneously activate the MAP kinase and the PI3K pathways, while also triggering calcium dependent peptidylarginine deiminase activity. It was this association of pathways that caused HERV activation, a phenomenon that was only part of more generally increased transcription of heterochromatic regions. The HERV transcripts were generally not polyadenylated. Some arose as a consequence of activation of HERV-based enhancers, while others were the result of unusually strong activation at some mostly transcription factor genes causing transcription to leak out of the HERV-free region that surrounds them. Altogether, our data emphasized the hazard associated with simultaneous activation of multiples signaling pathways by xenobiotics, while also providing a very general toolbox for the interpretation of HERV transcription.

## Introduction

Human endogenous retroviruses (HERVs) are vestiges of ancient retroviral infections of the germline. They are present in many more-or-less degenerated copies, classified in families depending on the tRNA used by the original retroviruses to prime the retrotranscription (H for histidine, W for tryptophane, etc.). Together, they represent between 5 and 15% of the human genome. Recently, HERVs have received much attention as next generation sequencing and CRISPR-Cas9 technology have provided evidence for a role of these sequences, together with other potentially transposable element (TE) such as LINEs and Alu sequences, in shaping transcriptional networks. Indeed, TEs are rich in regulatory elements initially necessary for the viral cycle and later possibly co-opted by the host, mostly in the context of immunity and antiviral defense.

Historically, HERVs have been also studied for their possible role in the onset of diseases. In particular, several autoimmune diseases including multiple sclerosis, type 1 diabetes mellitus, and rheumatoid arthritis have been associated with increased HERV activity, presence of virus-like particles in the blood, and production of antibodies against HERV-encoded proteins (Christensen 2010; Antony et al. 2011). While the role of these HERVs in the onset of autoimmune diseases has been extensively investigated, the mechanisms at the root of their increased transcription in the patients remain essentially unexplored.

Here, we have addressed this issue with a focus on HERVs frequently linked to multiple sclerosis (MS-HERVs). Multiple sclerosis (MS) is an immune-mediated disease of the central nervous system (CNS) associated with inflammation, demyelination, neuroaxonal damage, and neurodegeneration (Hafler 2004) (Lassmann 2018). While several lines of evidence indicate that MS is triggered by a combination of exogenous, environmental, and genetic factors, the molecular mechanisms at play are still poorly characterized, and MS remains an idiopathic and incurable disease afflicting hundreds of thousands of people worldwide.

In the first stages of the disease, damage to the CNS is mediated by inflammatory cell infiltrates, composed of CD8^+^ and CD4^+^ T cells, macrophages and activated local microglia (Lassmann 2018). CD4^+^ T cells have a pathogenic role by being autoreactive against myelin-derived antigens and by secreting pro-inflammatory cytokines (Dendrou et al. 2015). Inflammatory cytokines/chemokines are also significantly increased in serum of patients and proposed for monitoring disease activity. Thus, elevated levels of tumor necrosis factor-α (TNFα) and the chemokine CXCL8 (IL-8), both in cerebrospinal fluid (CSF) and serum, have been associated with disease progression and relapse episodes (Sharief, 1992; Hohnoki, 1998; Lund, 2004; Ishizu, 2005).

In earlier studies, we had shown that, in the absence of a pro-inflammatory stimulus, TNFα and IL-8 shared with HERVs a transcriptional repression machinery relying on histone H3 lysine 9 (H3K9) methylation and subsequent binding of HP1 proteins. In addition, we showed that in PBMCs from patients with MS, HP1-recruitment was reduced at both HERV sequences and at the promoter of TNFα. In a subset of patients, the inefficient HP1 recruitment could be further correlated with increased activity of the peptidylarginine deiminase PADI4. This calcium-dependent enzyme converts histone H3 arginine 8 (H3R8) into a citrulline and thereby destroys the HP1 binding site formed by methylated H3K9 (Sharma et al. 2012).

These observations pointed out the need for a more general understanding of the triggers and the chromatin events behind HERV transcriptional activity. To that end, we searched for a simple model to examine transcriptional activation of HERVs. In that context, we found that exposure of Jurkat T cells to the organochlorine pesticide dieldrin resulted in a reliable activation of several MS-HERVs. Examination of the signal transduction pathways at play provided much insight on the burden inflicted on the cell by environmental pollutant and revealed that it was the effectiveness of the chemical in activating several pathways beyond inflammation that provoked destabilizing of heterochromatin and transcription of HERVs. Genome-wide approaches further allowed us to detect several mechanisms leading to a HERV transcription readout. These may have a very general value in interpreting retroviral expression in patients with autoimmune diseases.

## RESULTS

### Dieldrin is an activator of several HERVs connected to MS

With the objective of identifying an activator of MS HERVs, we searched the literature for compounds potentially interfering with heterochromatin-mediated repression. The identified compounds were then tested in Jurkats, a T cell-derived tissue culture cell line for which we had previously shown activation of several MS HERVs by the combined effect of PMA and a calcium ionophore (Sharma et al. 2012). In addition, these cells are a well-characterized model for T cell activation (see for example (Brignall et al. 2017)). This activation was assessed by monitoring activity of the IL8 and TNFα, induced more rapidly than IL2.

A first good candidate for MS HERV activation was bpV(pic), an inhibitor of phosphotyrosine phosphatase previously shown to activate several HERV reporter constructs transfected into Jurkat cells (Toufaily et al. 2011). Consistent with these observation, bpV(pic) transiently stimulated transcription at some of the endogenous HERVs we monitored. This compound also caused solid activation of IL8 and TNFα (Figures S1A-S1D).

We next tested dieldrin, an organochlorine pesticide described as an inducer of histone acetylation in dopaminergic neuronal N27 cells (Song et al. 2010), while also promoting the production of IL8 in human neutrophils (Pelletier et al. 2001). Dieldrin was used in agriculture for the control of insect vectors of diseases. Although no longer used since the early 1990s, it is a global contaminant, listed under the Stockholm convention as a persistent organic pollutant (POP) (Convention 2009). Dieldrin has been associated with increased risk of breast cancer (Høyer et al. 1998) and Parkinson’s Disease (Kanthasamy et al. 2005). Titration experiments showed that Jurkat cells tolerated dieldrin at concentrations up to 100 µM for 1 hours with no significant cell death (less than 2% - Figure S1E).

All MS HERVs except HERV_K102 were induced 2-to 4-fold upon exposure to 100 µM Dieldrin for 30 min (Figure 1A). In comparison, activation of the Jurkat cells with 100 µM of the phorbol ester PMA stimulated IL8 and TNFα transcription less efficiently, while not inducing HERV expression at all (Figures 1B-1D). The concentration of PMA was here intentionally higher than that usually used (40 nM) to ascertain that the absence of HERV stimulation was not a consequence of insufficient dosage. At 100 µM, both dieldrin and PMA failed to activate TGFβ.

**Figure 1:**
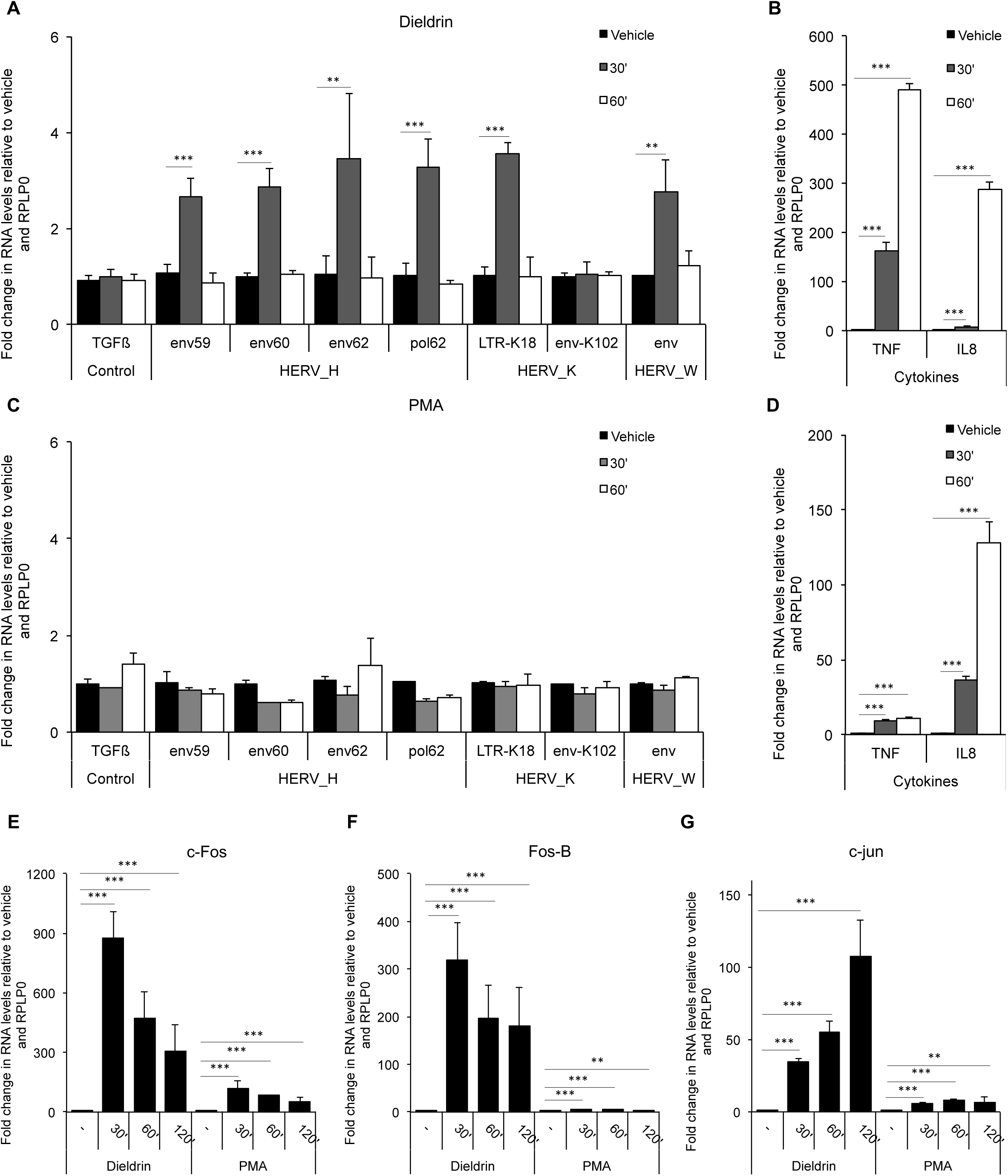
Dieldrin is an activator of several MS-HERVs. Jurkat T cells were treated with either DMSO (vehicle) or with 100µM of either Dieldrin or PMA for the indicated times. Abundance of mRNA for the indicated genes or HERVs was assessed by RT-qPCR. Data shown are means +/-SEM from three independent experiments. Significance (p-value) was estimated using the two-sided student t test.

The strong response of IL8 and TNFα to dieldrin prompted us to test additional PMA-responsive genes, and we observed a massive activation of immediate early genes c-Fos, Fos-B, and c-Jun (Figures 1E-1G). Examination of a series of genes markers for environmental, oxidative, osmotic, cytotoxic, hypoxic, endoplasmic reticulum, metal, or heat shock stress detected only HSP6A as upregulated more by dieldrin than by PMA (Figures S1F-S1G). In addition, we observed no DNA damage response as monitored by TRIM28 phosphorylation (Figure S1H).

We concluded from these observations that bpV(pic) and especially dieldrin are potent activators of T cells also inducing several MS HERVs. The lack of activation of these HERVs by PMA further indicated that their activity is not a systematic consequence of T cell activation. In the following sections, we will focus on dieldrin which was the most efficient at activating the HERVs, IL8, and TNFα, and also the more likely to be an issue for human health.

### Dieldrin provides the simultaneous activation of multiple pathways required for HERV transcription

To gain insight in the mechanisms allowing dieldrin to activate the HERVs, we next investigated the signal transduction pathways stimulated by dieldrin in comparison with PMA. A human phospho-kinase array confirmed that both PMA and dieldrin treatments resulted in the activation of the MAP kinase pathway as testified by the phosphorylation of the TXY motif in ERK1/2 proteins and of the SRF-kinases RSK1/2/3 (Figures 2A-2B, and S2AS2B). In that context, we examined phosphorylation of PKC, the primary target of PMA. Surprisingly, Dieldrin, unlike PMA, did not cause any visible change in PKC phosphorylation when examined by western blot with a pan phospho-PKC antibody (Figure 2C Phospho PKC, compare lanes 2 and 6), suggesting that dieldrin activates the MAP kinase pathway independently of the TCR/PKC pathway, possibly via tyrosine kinase receptors. To examine the impact of the ERK pathway on HERV transcription, we challenged dieldrin with two inhibitors of MEK (U0126 and PD98059). This resulted in decreased activation of HERVH env62 together with the AP1 genes, while not affecting TNFα (Figures 2D-2H and S2C-S2G).

**Figure 2:**
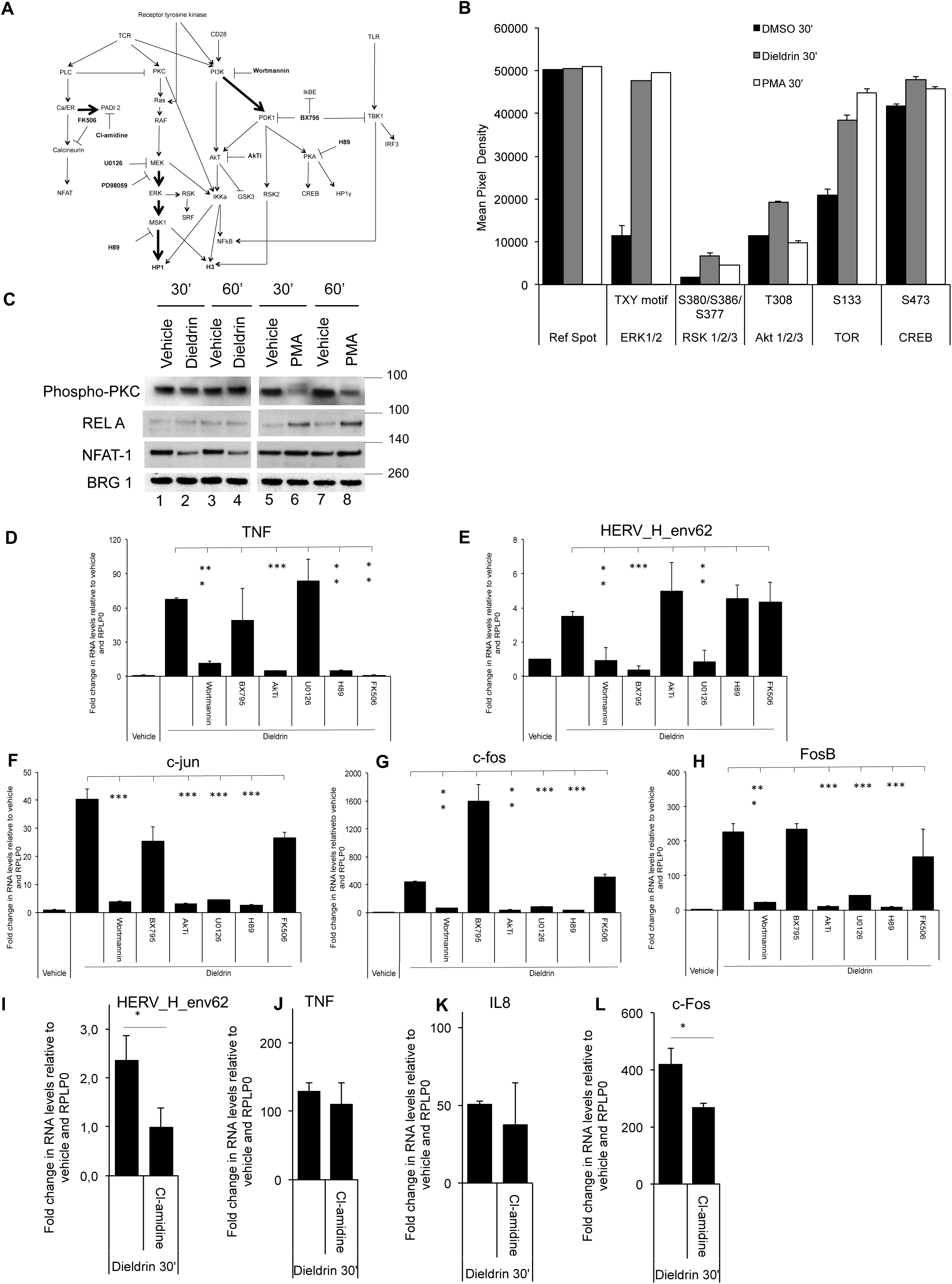
Dieldrin activates multiple signal transduction pathways. (A) Schematic of the signal transduction pathways under scrutiny. Used inhibitors are indicated. Transitions demonstrated as relevant for HERV H env62 transcription are indicated with bold arrows. (B) A human phospho-kinase antibody array was probed with extracts from Jurkat cells treated with either DMSO, dieldrin, or PMA for 30 min. as indicated. The array processed as a western and revealed by ECL. Graphic shows selected antibodies (average from 2 spots). The full experiment is shown S2A and S2B. (C) Jurkat cells were treated with either DMSO (vehicle), dieldrin, or PMA for the indicated times. Cell extracts were resolved by PAGE and western blotting was carried out with the indicated antibodies. Nearest size marker is indicated to the right (Kb). (D-L) Jurkat cells were treated for 30 min. with either DMSO (vehicle) or with 100µM of Dieldrin in the absence or in the presence of the indicated small molecule inhibitors. Abundance of mRNA for the indicated genes or HERVs was assessed by RT-qPCR. Data shown are means +/-SEM from three independent experiments. Significance (p-value) was estimated using the two-sided student t test.

Jurkat cells have elevated basal activity of the PI3K/AKT pathway due to a mutation in the PTEN gene. This activity accounted for the high levels of AKT S473 phosphorylation detected on the phospho-kinase arrays (Figure S2B). Nevertheless, both PMA and dieldrin caused increased TOR phosphorylation at S2448, suggestive of a further activation of the pathway (Figures 2B). The activity of the PI3K/AKT pathway was necessary for the activation by dieldrin of all the genes we monitored, including HERVH env62, as demonstrated by the effect of the PI3K inhibitor wortmanin (Figures 2D-2H). We noticed however that, unlike PMA, dieldrin did not induce detectable REL A phosphorylation (Figure 2C REL A, compare lanes 2 and 6), suggestive of an atypical kinase cascade. To investigate this further, we followed up on the fact that dieldrin but not PMA caused phosphorylation of Akt at T308 (Figures 2B), a residue modified by PDK1 (Manning and Toker 2017). Inhibition of this kinase by BX795 resulted in repression of HERVH env62, without affecting any of the other genes. As BX795 also targets TBK1 and IKBKE, it is difficult to draw definitive conclusions on the pathway at play, but this molecule may have a potential therapeutic value in diseases associated with increased HERV expression.

Finally, we examined pathways activated by calcium influx. Both dieldrin and PMA caused phosphorylation of CREB at S133, a modification catalyzed either by PKA or by the CamK in the presence of Ca2+. The use of the PKA-inhibitor H89 proved however that this pathway, although important for TNFα and AP1 activity, did not impact on HERVH env62 (Figures 2D-2H). In parallel, we found that dieldrin but not PMA caused NFATc2 de-phosphorylation, an indicator of the activation of the calcium/calmodulin pathway (Figure 2C NFAT-1, compare lanes 2 and 6). Consistent with this, the Calcineurin inhibitor FK506 interfered with activation of TNFα by dieldrin, however without affecting neither the AP1 genes, nor HERVH env62 (Figures 2D-2H). This conflicted with our earlier observation that PMA should be combined with a calcium ionophore to activate HERVs (Sharma et al. 2012). This prompted us to envision a third route for activation by calcium and a possible implication of peptidylarginine deiminases. These enzymes are highly dependent on intracellular Ca+ concentrations (Arita et al. 2006), and we have previously shown that PADI4 is a positive regulator of several HERVs (Sharma et al. 2012). The PADI inhibitor cl-amidine caused a 2-fold reduction in the effect of dieldrin on HERV H env62, had a small but significant effect of activation on c-Fos, while not affecting activation of TNFα and IL8 (Figures 2I-2L).

Altogether, this series experiments suggest that the specificity of dieldrin resides in its ability to simultaneously activate multiple pathways, whereof the MAP kinase and the PI3K/Akt together with calcium influx are essential for HERV activation. Our data further suggest that the effect of calcium on both HERVs and AP1 genes is mediated by PADI enzymes rather than calcineurin and NFAT. This is consistent with our earlier observations showing that citrullination of histone H3 at arginine 8 relieves HP1-mediated repression at MS HERVs (Sharma et al. 2012).

### Dieldrin interferes with H3K9me3/HP1-mediated silencing

To investigate the role played by the H3K9me3/HP1 axis in the effect of dieldrin on MS HERV activity, we first probed the silencing machineries keeping these HERVs in check in the Jurkat cells. To that end, cells were exposed for 16h to either chaetocin, an inhibitor of the histone H9K9 tri-methylase SUV39H, BIX-01294, an inhibitor of EHMT2/G9a, a methyltransferase that mono- and dimethylates H3K9, or azacytidine, a nucleotide analog that inhibits the maintenance DNA methylase DNMT1. In these experiments, only chaetocin treatment caused reactivation of all the HERVs examined in Figure 1, strongly suggesting that these retroviral sequences are silenced mainly by H3K9 tri-methylation and SUV39H proteins, rather than DNA methylation (Figures 3A, S3A, and S3C). Azacytidine had a strong effect on other heterochromatic loci as documented by a strong induction of the imprinted gene H19 (Figure S3D). At note, chaetocin also activated the pro-inflammatory cytokine TNFα and IL8 more efficiently than azacytidine (Figures 3B, and S3E), sustaining the role played by the H3K9me3/HP1 axis in controlling inflammation.

**Figure 3:**
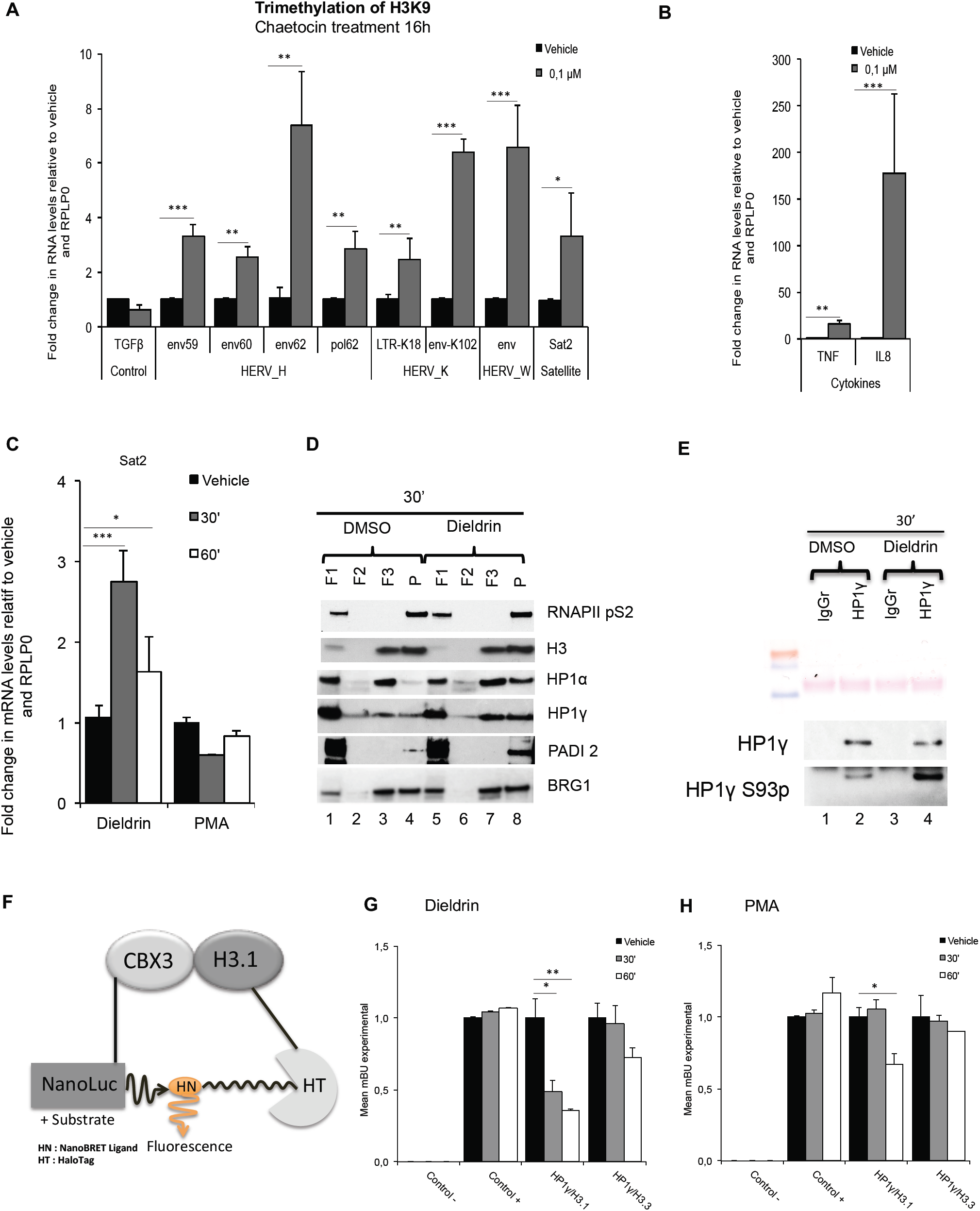
Dieldrin interferes with H3K9me3 and HP1-mediated transcriptional silencing. (A-B) Jurkat cells were treated for 16 hours with either DMSO (vehicle) or with 0.1 µM of the SUV39H-inhibitor chaetocin as indicated. Abundance of mRNA for the indicated genes or HERVs was assessed by RT-qPCR. (C) Jurkat cells were treated for 30 min. with either DMSO (vehicle) or with 100µM of Dieldrin. Abundance of Sat 2 RNA was assessed by RT-qPCR. In panels A-C, data shown are means +/-SEM from three independent experiments. Significance (p-value) was estimated using the two-sided student t test. (D) Jurkat cells treated as in C were subject to fractionation as indicated in the Material and Method section to distinguish a fraction with mostly RNAPII S2p (F1), one with mostly histone H3 (F3), and one with both RNAPII S2p and histone H3 (P). Fractions were resolved by PAGE, and wersten blots were carried out with the indicated antibodies. (E) Extracts were prepared as in C, then used for immunoprecipitation with either mouse IgGs or anti-HP1γ antibodies. Western blot was revealed with the indicated antibodies. (F) Schematic of NanoBret. When NanoLuc-Cbx3 and H3.1-HaloTag fusion proteins are in proximity, energy produced by luciferase is transferred to the HaloTag, resulting in a shift of wavelength. (G-H) HEK293 were transfected with plasmids expressing the constructs described in F. 36 hours later, the cells were treated with DMSO (vehicle), Dieldrin, or PMA for the indicated times. The shifted wavelength was then measured with a luminometer. Dta shown are means +/-SEM from three independent experiments. Significance (p-value) was estimated using the two-sided student t test.

We next investigated whether dieldrin would have an effect on transcription of heterochromatinized sequences other than the HERVs. In particular, we examined expression of Sat2 repeats localizing mainly to heterochromatic regions on chromosomes 1 and 16, and regulated by SUV39H and H3K9me3 (see for example (Wang et al. 2013) and Figure 3A last histogram). The primer set we used detected a site on chromosome 16 that we found stimulated by dieldrin, but not by PMA (Figure 3C). This observation prompted us to determine whether dieldrin had a global effect on heterochromatin. Immunofluorescent staining of HP1γ and staining of the DNA with DAPI did not reveal any overt disassembly of heterochromatic foci (Figure S1H). Cell fractionation experiments allowed for a more detailed analysis. Using a previously described protocol (Reyes et al. 1997), we isolated a very soluble fraction rich in RNAPII and poor in chromatin (as determined by histone H3 content – Figure 3D Fraction F1), a fraction rich in chromatin but containing no RNAPII (F3, regarded as containing inactive chromatin), and a pellet containing both chromatin and RNAPII (P, regarded as transcribed chromatin). Upon exposure to dieldrin, the very heterochromatic HP1α and more euchromatic HP1γ became enriched in transcribed chromatin, suggesting commitment of these proteins in transcription (Figure 3D, compare lanes 4 and 8). A commitment of HP1γ in transcription was compatible with its phosphorylation at S93 (Figure 3E, compare lanes 2 and 4), a mark of euchromatic HP1γ (Lomberk et al. 2006). In the fractionation experiment, HP1γ surprisingly also showed increased binding to inactive chromatin, possibly preparing for rapid re-compaction of these regions or participating in transcription. We finally examined the fractions for PADI2, which is the only PADI enzyme expressed in Jurkat cells. Western blots revealed an increased accumulation of this enzyme in transcribed chromatin in cells exposed to dieldrin (Figure 3D PADI2, compare lanes 4 and 8).

Finally, we investigate directly the effect of dieldrin and PMA on interaction between histone H3 and HP1γ. To that end, we used NanoBret, a bioluminiscence resonance energy transfer-based technic that allows quantifying protein-protein interactions in live cells (Figure 3F). Plasmids encoding NanoLuc-HP1γ or H3-HaloTag fusion proteins were co-transfected into HEK293T cells. In these assays, dieldrin but not PMA specifically reduced interaction between HP1γ and H3.1, while not affecting interaction with H3.3 (Figure 3G and 3H). This was consistent with H3.1 being the more H3K9-methylated histone H3 variant, while H3.3 is loaded mainly at actively transcribed sites (Loyola et al. 2006).

Together, these data suggest that the transcriptional activation of MS HERVs by dieldrin involves a weakening of chromatin-dependent silencing mediated by H3K9me3 and HP1 proteins that also affects other heterochromatic sequences. The reduced HP1-H3 interaction may possibly be linked to the activation calcium influx and PADI2-mediated citrullination of histone H3, combined with the HP1γ phosphorylation, previously shown to be a consequence of activation of the MAPK pathway (Harouz et al. 2014).

### Dieldrin causes extensive transcriptional disruption

To gain insight on the global effect of dieldrin on gene expression, we used next generation sequencing of a poly(A)-enriched random-primed cDNA library to analyze the transcriptome after 30 min of exposure. The transcriptome was compared to that of untreated cells or activated with PMA.

Principal component analysis clearly positioned the effect of dieldrin away from that of PMA (Figure 4A). We identified a total of 246 genes significantly (pVal<0.05) upregulated by dieldrin, with strong activation of JUN/FOS, EGR1/2, CD69, and DUSP1, indicative of a strong mitogenic effect of the molecule. We also noted a more than 5-fold activation of the SERPINE1 gene, encoding plasminogen activator inhibitor type-1 (PAI-1) previously shown to favor survival of Jurkat cells (Placencio et al. 2015). Several lines of evidence confirmed that dieldrin was a potent activator of the MAPK pathway, while PMA was more efficient at activating NF-κB signaling. First, a KEGG pathway analysis identified MAPK signaling as the pathway most enriched in dieldrin-inducible genes, while PMA-induced genes mapped better to NF-κB than to MAPK signaling (compare Figures 4C and 4D). Next, among the 56 genes activated by both dieldrin and PMA, MAPK responsive genes clustered among genes activated more by dieldrin than by PMA (yellow genes are mostly in the red zone, Figure 4E). Finally, examination of ENCODE ChIP-seq data identified STAT3, TCF3 and RELA peaks as the most frequently observed in the vicinity of dieldrin-induced genes. The same transcription factors were significantly enriched in the vicinity of PMA-induced genes, but with a higher score for the NF-κB p65/RELA. At note, CD83 and TNFAIP3, two NF-κB responsive genes involved in the dampening of the inflammatory reaction were activated more by PMA than by dieldrin, suggesting a less contained effect of diedrin (Figure S4A-S4B, and (Giordano et al. 2014; Reinwald et al. 2008)). Finally, the transcriptome analysis confirmed that dieldrin activated IL8 and TNFα, while also revealing that these are the only cytokine genes responding to the pesticide in 30 min. (Table S1).

**Figure 4:**
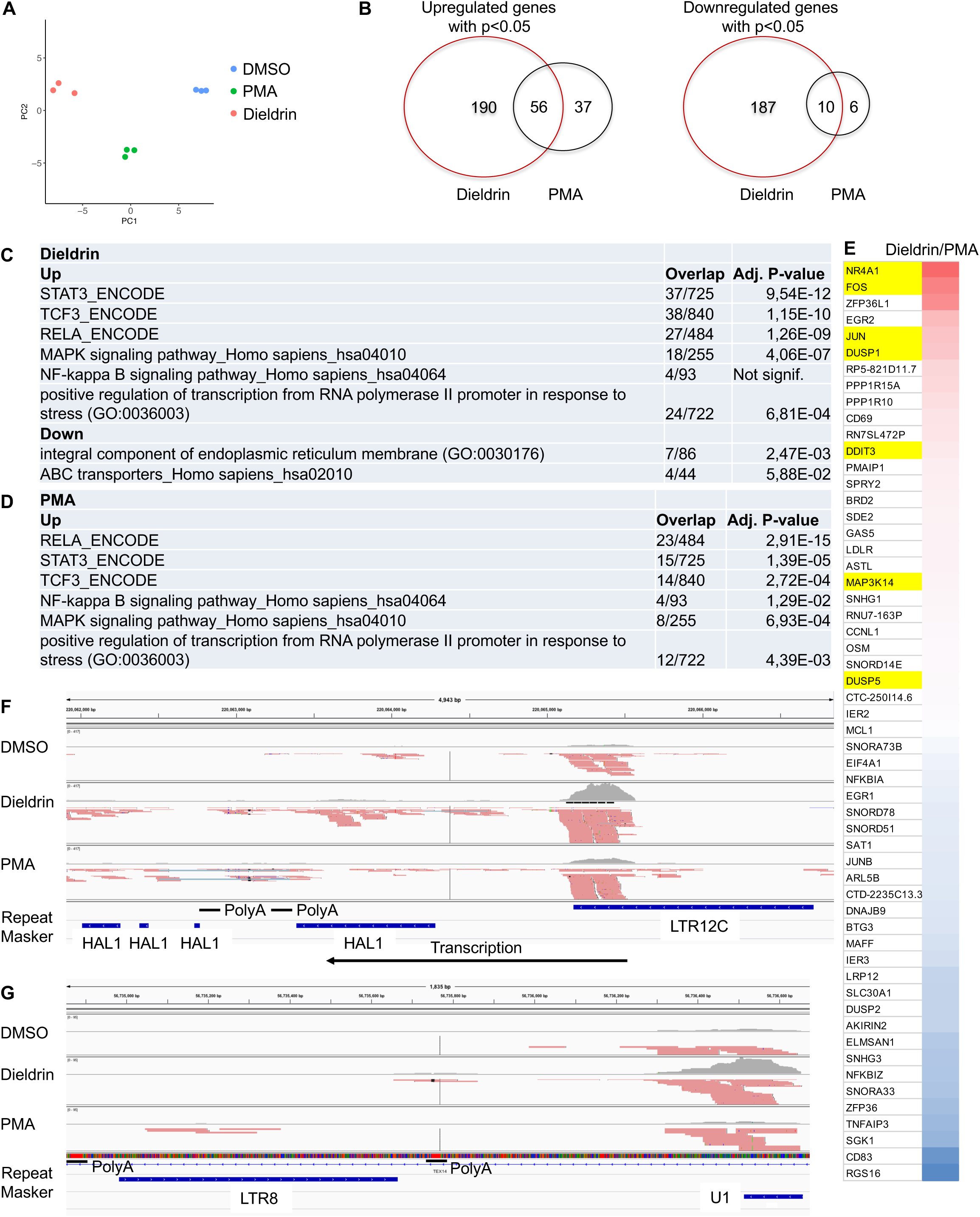
Transcriptome analysis of Jurkat cells exposed to dieldrin for 30 min. (A) Principle component analysis on the 485 genes the most affected by DMSO, PMA, or Dieldrin. (B) Venn diagrams reporting genes up- or down-regulated dieldrin or PMA as compared to DMSO. (C) List of genes upregulated both by dieldrin and by PMA. Scale indicates the dieldrin/PMA ratio. Dark red indicates strong activation by dieldrin, dark blue, strong activation by PMA. Yellow highlights indicate genes listed as regulated by MAPK in the KEGG pathway hsa04010. (D-E) Gene ontology analysis of RNA-seq data. (F) Screen shot from IGV genome browser showing transcripts initiated in a HERV and penetrating a region rich in A/T tracts. (G) Screen shot from IGV genome browser showing U1 transcripts captured in the poly(A) enrichment most likely due to to the proximity of HERV-born poly(A/T) tracts.

Interestingly, dieldrin also caused moderate (less than 2-fold) but significant (p<0.05) transcriptional repression of a large set of genes (197 genes – Figure 4B). This is reminiscent of the extensive transcriptional repression induced by heat shocks in mammalian cells (Mahat et al. 2016). In this context, we noted the moderate (less than 2-fold) activation of heat shock proteins HSPA6, HSPA1A/B, and HSPH1, suggesting some similarities between heat shock and dieldrin-induced stress (Table S1). The down-regulated genes were of very diverse function but included genes encoding components of the endoplasmic reticulum (ER) and ABC transporters. However, ER-stress markers DDIT3/CHOP and DNAJB9 were not activated more by dieldrin than by PMA (Table S1 and Figures S1F-S1G). In addition, we did not detect any cleavage of ATF6, a hallmark of ER stress (data not shown). Possibly, the ER stress is very moderate or in an early phase.

We next counted reads mapping inside regions annotated as “LTR” in RepreatMasker (A.F.A. Smit, R. Hubley & P. Green RepeatMasker at http://repeatmasker.org). This mapping was facilitated by the long (150 nucleotides) paired-end reads of our transcriptome analysis. Yet, this approach detected surprisingly few retroviral sequences activated by dieldrin, thereby contrasting with the PCR data. However, examination of some of the few up-regulated LTRs revealed that the reads mapped to transcripts that had been retained in the poly(A) selection not because of post-transcriptional polyadenylation, but because they were encoded by DNA rich in tracts of Ts. In the example shown Figure 4F, the LTR12C sequence immediately neighbors several LINE-1 retrotransposons of the “half-L1” (HAL1) category. The region covered by these LINEs is composed of 31% Ts, including a tract of 24 Ts and one of 16 Ts. LINEs are also known as poly(A) retrotransposons because they contain a poly(A) tract required for their retrotransposition (Doucet et al. 2015). From these observations we concluded that the HERV transcripts mostly do not get polyadenylated. Genome-encoded T tracts were also likely to account for the detection of the non-polyadenylated U1 small nuclear RNA at least one RNU1 gene (Figure 4G). This attracted our attention as polyadenylated U1 had been detected in patients with MS (Spurlock et al. 2015).

### HERVs are transcribed as enhancers or as a consequence of gene “spillover”

To evaluate the overall impact of dieldrin on transcription, we repeated the RNA-seq of cells stimulated or not by dieldrin using a cDNA library constructed without poly(A) enrichment. To estimate the function of the genomic regions the more affected by dieldrin, we assigned the reads to the different chromatin states as defined by the 15-state model of the NIH Roadmap Epigenomics Mapping Consortium. As the epigenome of Jurkat cells was not available, we used and averaged 6 different epigenomes from T cells and that of the human myelogenous leukemia cell line K562. Variations in % of each state is shown as a boxplot in Figure 5A. The analysis shows an overall reduced activity at transcription start sites (TssA and TssBiv). This may be a reflection of the broad downregulation of genes observed in the poly(A) selected transcriptome data. In contrast, regions annotated as enhancers showed increased transcription, a likely consequence of the activation of the mitogenic response. Regions annotated as quiescent, containing no or few histone modifications and enriched in silent genes, and to a lesser extend regions annotated as Heterochromatin, Repeats, or Weak repression by Polycomb also showed increase in transcription, suggestive of a reprogramming of heterochromatin, a phenomenon that may include the weakened H3K9me3/HP1-mediated silencing described in Figure 3.

**Figure 5:**
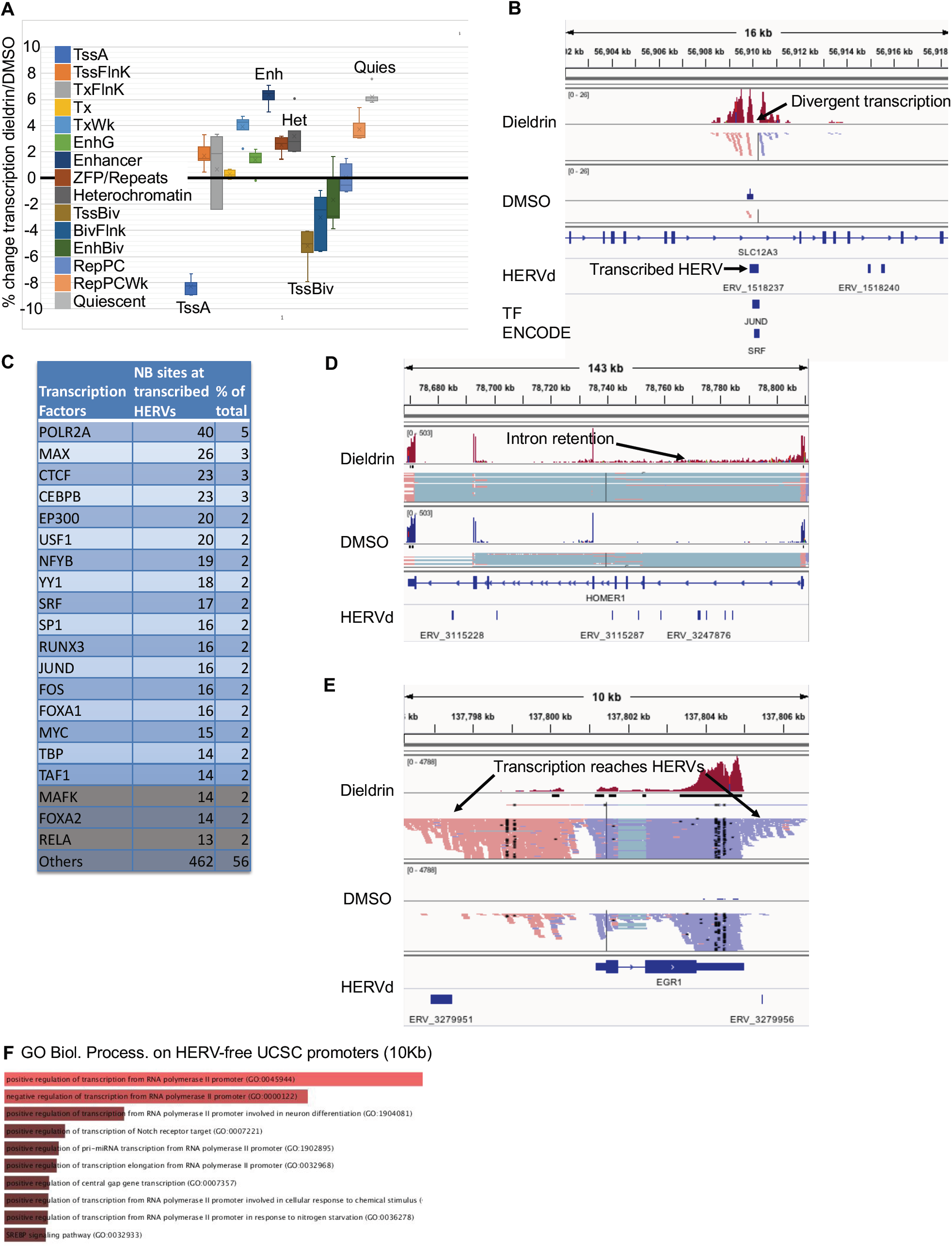
HERV transcripts originate from enhancers and from transcription of gene neighborhoods. (A) Quantification of reads mapping inside chromatin states defined by the “Epigenome Road Map Consortium” 15 states model using RNA-seq data obtained from Jurkat cells treated for 30 min. with either DMSO (vehicle) or dieldrin. Values shown are % variations in number of reads (dieldrin vs. DMSO) in each chromatin state adjusted by the total number of reads in each RNA-seq data set, averaged from quantification of 4 epigenomes from primary T cells (E039, E041, E043, E044) and from that of the K562 leukemia cell line (E123). (B) Example of a HERV becoming a site of divergent transcription upon exposure of the Jurkat cells to dieldrin (C) List of transcription factions with peaks in the ENCODE data mapping to HERVs showing divergent transcription activated two-fold or more in the presence of dieldrin. Each transcription factor is associated with the number of transcribed HERVs where their peaks are found. (D) Example reads covering HERVs but due to poor processing of an mRNA. (E) Example of dieldrin-induced transcription bleeding outside of the HERV-free region. (F) Gene ontology analysis of genes having no HERVs in the 10Kb region located upstream of their transcription start site (TSS).

We next counted reads mapping inside sequences of retroviral origin. In this approach, we preferred the HERVd library of HERVs, where RepeatMasker sequences likely to derive from a same insertion event are annotated as a single HERV (see example Figure S5A and https://herv.img.cas.cz/). This approach detected hundreds of HERV sequences activated by dieldrin. To analyze these sequences, we first focused on sites of divergent transcription. Divergent transcription is a hallmark of promoters and enhancers and the gap between the sense and the reverse transcripts frequently designate sites of transcription factor binding (see example of the FOS gene in Figure S5B and (Andersson et al. 2015)). Examples of HERV sequences hosting divergent transcription are shown Figure 5B, S5C and S5D. When examining globally genes located in neighborhood of these HERVs using GREAT (http://bejerano.stanford.edu/great/public/html/index.php), we observed an enrichment in genes involved in T-cell activation and negative regulation of apoptosis (Figure S5E). This strongly suggested that HERV sequences have been coopted by the mammalian cells for cell survival and activation of the T-cell immune response.

We next examined sites of divergent transcription within HERVs for overlap with sites of transcription factor binding from the ENCODE consortium. This allowed to estimate which transcription factors were likely to use HERVs as binding sites upon stimulation by dieldrin. The approach designated classical players of T cell activation, including MYC/MAX, SRF, and FOS. Peaks of CTCF, p300, and CEBP further supported an enhancer function for these HERVs (Figure 5C).

While sites of divergent transcription illustrated the role of HERVs in transcriptional activation, we also noted that many HERV sequences were transcribed as a consequence of the transcription of overlapping or nearby genes. In some cases, reads aligning to HERVs were detected because of altered pre-mRNA maturation. This was for example noted at the HOMER1 gene where reads covering the six first introns were abundantly detected, possibly illustrating an effect of dieldrin on mRNA processing (Figure 5D). In addition, the strong activation of immediate early genes occasionally resulted in transcription of HERV sequences either because of the very abundant promoter RNAs or defective transcription termination (see an example Figure 5E). This later observation pointed out the risk of excessive gene activity, and prompted us to investigate which genes were more likely to be in “HERV-free” neighborhoods. This approach very clearly designated transcription factors (Figure 5F) and examination of for example the MYC gene clearly revealed a relatively large region mostly upstream of the gene and devoid of HERV sequences (Figure S5F). This was confirmed by examining HERV “density” (number of HERV sequences inside the coding region divided by the length of the coding region), an approached showing that almost 80% of transcription factors have a HERV density lower than average (Figure S5G). These observations suggest that highly inducible genes are buffered in HERV free regions to accommodate for promoter transcription and inefficient transcription termination, but that in the case of dieldrin exposure, this buffering is not sufficient.

## Discussion

Here, we wished to explore to what extend transcriptional activity at HERVs is a systematic consequence of the cellular stress response and whether that could explain HERV activity as it is observed in several autoimmune diseases. Activation of HERVs by stress has been envisioned extensively, but always with working models involving protein synthesis from the HERVs, and subsequent inflammatory or immune response. To investigate a possible cell-autonomous link between HERV transcription and stress, we identified an environmental pollutant able to induce a transient burst of transcription at several of HERVs in a cell line frequently used to study mitogenic signaling. First, we examined which signaling pathways allowed dieldrin to activate HERV transcription. We found that a characteristic of dieldrin was its capacity to activate multiple pathways, including the MAP kinase, the PI3K/AKT, the calcium/calcineurin, and possibly the cAMP/PKA pathways. It seemed that this pleiotropy is at the origin of the effect of dieldrin on the HERVs. We knew from earlier experiments that PMA associated with an ionophore allowed activation of HERVs (Sharma et al. 2012). We discovered here that it is the combine effect of MAPK/PI3K activation with calcium influx that allows for HERV transcription, and while PMA alone induces only MAPK/PI3K activation, dieldrin combines activation of these pathways with calcium influx. It is noteworthy here that in Jurkat cells, the basal level of the PI3K pathway is high, because of a mutation in the PTEN gene. For that reason, these cells may have been a particularly favorable model to examine HERV transcription. The role of the PI3K pathway in the transcriptional activity of HERVs and the possibility of specifically inhibiting HERVs with PDK1 inhibitors have been serendipitous observations that may bring renewed interest for studies linking the PI3K pathway to inflammatory disorders (see for example (Xie et al. 2014)).

Interestingly, HERV transcription, while requiring calcium, did not require calcineurin activity. This prompted us to examine PADI activity that we had previously found activated in patients with MS (Sharma et al. 2012). We found that dieldrin caused PADI2 to associate with chromatin, and that the PADI-inhibitor cl-amidine interfered with transcriptional activation of HERV_H-env62 and of the FOS gene. H3 citrullination by PADI enzymes disrupts HP1 binding to the H3K9me3 mark, consistent with the NanoBret experiments showing a decreased interaction of histone H3 with HP1γ in the presence of Dieldrin. The H3 citrullination is therefore expected to interfere with heterochromatin integrity and promote re-localization of the HP1 proteins and subsequent transcriptional activation. This mechanism may play a hitherto overlooked role in chromatin reprogramming in response to stress. It may be part of the re-localization of genes and gene-interactions recently associated with T-cell activation (Robson et al. 2017). It is here of interest to note that PADI enzymes are also involved in more dramatic chromatin destructuring in neutrophils in the form of neutrophil extracellular traps (NETs) designed to ensnare and kill extracellular bacteria (Rohrbach et al. 2012). In addition, the re-localization of HP1 proteins by H3 citrullination presents several similarities to that induced by the Janus kinase 2 (JAK2). This kinase, that is frequently activated in hematological malignancies and up-regulated in patients with MS (Conti et al. 2012), interferes with HP1α binding to chromatin by phosphorylating H3Y41 (Dawson et al. 2009). In *Drosophila*, the higher levels of heterochromatin resulting from decreased levels of activated JAK are associated with suppression of hematopoietic tumor-like masses, increased resistance to DNA damage, and longer lifespan (Silver-Morse and Li 2013).

Examination of the transcriptome data from the poly(A)-enriched libraries confirmed that dieldrin is a strong activator of the MAPK pathway and revealed that HERV transcripts activated by dieldrin are mostly not polyadenylated. The poly(A) tracts found associated with transcripts were all encoded by the genome and, in the cases we examined, belonged to neighboring LINE or Alu sequences. In addition, the human genome contains several HERVs followed by poly(A) tails that are in fact LINE-processed pseudogenes from HERV mRNAs (Pavlícek et al. 2002). These pseudogenes are remarkably numerous in the HERV-W family, that is one of the most frequently associated with MS (see for example (Gjelstrup et al. 2018)). It is therefore tempting to speculate that poly(A) tracts, scars of LINE/SINE retro-transposition events, provide increased stability to some HERV-encoding transcripts and possibly allow for their occasional translation. This would be yet an additional example of synergy between HERVs and retrotransposons.

The analysis of the RNA-seq data from non-poly(A)-enriched libraries allowed us to identify numerous sites of divergent transcription inside HERV sequences. Combined with the absence of polyadenylation, this strongly suggest that a fraction of HERV transcripts induced by dieldrin are enhancer RNAs (eRNAs). We noted also that the very rapid and transient transcriptional activation of HERVs is very similar to the transcriptional behavior of enhancers (Zaret et al. 2008). A co-optation of HERV sequences as enhancers has been suggested earlier (Frank and Feschotte 2017; Chuong et al. 2013). In this context, the RNA-seq data further showed that, in response to stress or mitogens, HERVs provide binding sites for essential transcription factors such as MYC/MAX, AP1, SRF, and NFk-B in proximity to genes regulating T cell differentiation. Beyond the HERVs, examination of changes in transcriptional activity within the different annotated chromatin states designated enhancers as extensively reacting to dieldrin. Interestingly, poised enhancers frequently carry H3K9me3-marks (Zentner et al. 2011), and it is possible that part of the PADI-driven citrullination and the HP1-mobilization may happen at actived enhancers. This would explain why the FOS gene is sensitive to PADI-inhibition by cl-amidine, while it is activated by chaetocin (Figure S3F).

The RNA-seq data from non-poly(A)-enriched libraries revealed that HERVs also seems to get “accidentally” activated by dieldrin, either because they are located inside the coding region of genes subject to intron retention, or because they are located in proximity to enhancers, or promoters, or downstream of improperly terminated genes. This suggested a toxicity of dieldrin also reaching the transcription and the RNA-maturation machineries. In addition, this phenomenon unveiled that genes encoding transcription factors are frequently located in HERV-free zones with a buffering region of 10Kb in average upstream of the TSS and downstream of the TTS. This buffering region seems sufficient to avoid transcription of HERVs in the unstimulated cells while being surged over upon exposure of the cells to dieldrin. At the promoter of transcription factors strongly induced by dieldrin (such as JUN, FOS, EGR1 …), the combined activation of several signal transduction pathways may participate in generating promoter RNAs reaching beyond the buffer region. Downstream of the coding region, R-loop–mediated H3K9me marks have been implicated in slowing down the RNA polymerase II (RNAPII) at the end of some genes, allowing 5’-3’ exonuclease Xrn2 to catch up with the RNAPII and trigger its release from the DNA template (Skourti-Stathaki et al. 2014). The reduced binding of HP1 proteins to histone H3 induced by dieldrin may interfere with this mechanism and favor transcription of downstream HERVs. Such an increase in transcription outside of genes may be considered in the perspective of one of the few transcriptome studies performed on patients with MS and that concluded that only few genes were deregulated while RNA misprocessing was extensive. This misprocessing included a mysterious polyadenylation of U1, a phenomenon we also observe with dieldrin, and that we now can explain by transcription reaching into A tracts.

Together, these observations prompt us to propose a general model for HERV transcription that combines activation of “enhancer HERVs” and excessive transcription both upstream and downstream of genes as a consequence of unusual stimulation and imperfect transcript termination (Figure 6). Dieldrin has been an interesting tool to uncover these transcriptional behaviors, although, at this point, we have no evidence to incriminate dieldrin in the onset of MS. Yet, we believe that environmental pollutants simultaneously stimulating transcription, while also interfering with termination and silencing machineries are good candidates to be examined for a better understanding of MS and other autoimmune diseases involving transcription of HERVs.

**Figure 6:**
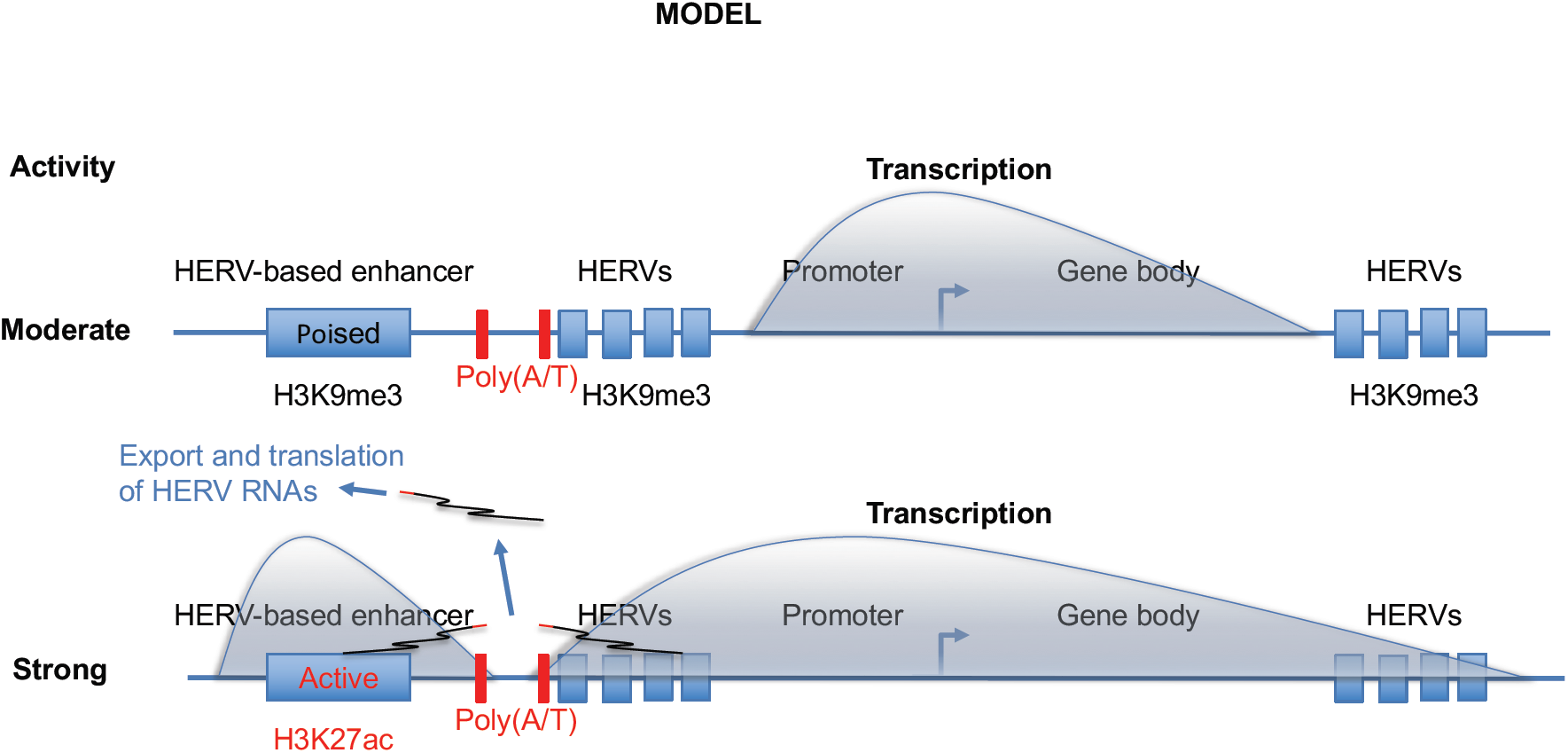
Model. (1) Physiological stimulation of gene expression results in transcriptional activity contained within a HERV-free region framing many genes (mostly encoding transcription factors). Strong multi-pathway stimulation for example by xenobiotics like dieldrin results in transcription beyond the borders of the HERV-free region. This effect may be amplified by the eventual interference of the xenobiotic with transcription termination and RNA processing. (2) Transcription through the HERVs may disturb the silenced chromatin they contain. In parallel, physiological and non-physiological activation of poised enhancers organized in silenced chromatin may also participate in restructured heterochromatin. When these enhancers are HERV-based, this will be an additional source of HERV transcripts. (3) HERV transcripts are not polyadenylated post-transcriptionally, but poly(A/T) tracts present in the genome and originating from A/T rich retrotransposons occasionally allow stabilization, export, and translation of the transcripts.

## Material and methods

### Antibodies, and chemicals

Antibodies used for Western-Blot: anti-Phospho-PKC (Cell signaling 9371T), anti-REL A (Cell signaling 3033T), anti-NFAT-1 (BD Transduction Laboratories 610702), anti-BRG1 (Euromedex 2E12), for cell fractionation experiments: anti-RNA Pol 2 pS2 (Abcam 5095), anti-H3 (Abcam 1791), anti-HP1α (Euromedex, Clone 2G9), anti-HP1γ (Euromedex, Clone 1G6), anti-PADI2 (Abcam 56928) and anti-BRG1 (Euromedex 2E12). Chemicals used in this study: Dieldrin and PMA (used on Jurkat and HEK293) and bpV(pic) were purchased from SIGMA (Ref 291218, P1585 and SML0885 respectively). The inhibitors used for the pathways determination: wortmannin (Sigma W3144), BX795 (Sigma SML0694), Akt1/2 kinase inhibitor (Sigma A6730), H89 (Sigma B1427), U0126 (EMD Millipore 19147), FK506 (Enzo Life Sciences ALX 380 008) and PD98059 (Selleckchem S1177). 5-Aza-2’-deoxycytidine, chaetocin and BIX-01294 were purchased from Sigma, A3656, C9492 and B9311 respectively. Cl-amidine are from EMD Millipore reference 506282.

### Cell culture

HEK293 cells were cultured in Dulbecco’s modified Eagle’s medium (DMEM, Gibco BRL), and Jurkat cells were cultured in RPMI-1640, all with 10% decomplemented fetal bovine serum (FBS) and 100 U.ml-1 penicillin-streptomycin at 37°C in a 5% CO2 incubator. Jurkat cells were treated with phorbol myristate acetate (PMA) or dieldrin at 100 µM, with bpV(pic) 10µM for the indicated times, with wortmannin (10mM), BX795 (6µM), Akt1/2 kinase inhibitor (40µM), H89 (40µM), U0126 (12µM), FK506 (100nM) and PD98059 (25µM) for 1h and with 200 µM Cl-amidine for exactly 16 h in complete cell culture medium. 5-Aza-2’-deoxycytidine, chaetocin and BIX-01294 were used at the indicated concentrations for 16h.

### mRNA, protein quantification, cell fractionation

Total RNA from Jurkat cells was extracted with a phenol-chloroform-based method and quantified with an ND-1000 (Nanodrop). After DNase treatment (Roche), reverse transcription was performed using SuperScript III (Invitrogen) and random hexanucleotides according to the manufacturer’s instructions. Complementary DNAs were quantified by RT-qPCR (Mx3005P, Stratagene) using SYBR Green PCR master mix (Applied Biosystems). PCR primers are available upon request. Data are shown relative to RPLP0 considered invariant. Total proteins were separated by electrophoresis on 4–12% gradient PAGE gels (Bio-Rad, #345–0124) and transferred on nitrocellulose membrane (Bio-Rad, #1620115) for western blot with indicated antibodies. Cell fractionation was carried out as previously described (Reyes et al. 1997).

### NanoBRET

NanoBRET kit from Promega was used according to the manufacter’s instructions. Briefly, in HEK293T, we carried out transient transfections in a 6-well plate using FuGENE HD (Promega). For each well, we incubated 2 µg of H3.1 HaloTag fusion vectors (or H3.3 HaloTag or control) and 0,002 µg of CBX3-NanoLuc for 10 min at room temperature with a mix of 1 μl of FuGENE and OptiMEM up to 100 μl. We then incubated cells (8×10^5^ in 2mL per well) in DMEM supplemented with 10% FCS with the final DNA-FuGENE mix (50 μl/well). 24h post-transfections, the cells were re-plated into 96 well plates (2.2×10^5^ cells / mL), and fluorescent NanoBRET HaloTag^®^ 618 Ligand (for experimental samples) and control samples (no fluorescent ligand) were added. After 24h, we carried out assays by treated cells with our compound (dieldrin or PMA) for 30 min, 60 min and then incubated them with the NanoBRET NanoLuc^®^ Substrate, and donor and acceptor signals are measured on an instrument capable of measuring dual-filtered luminescence equipped with appropriate filters. The corrected NanoBRET ratio is calculated, which is a subtraction of NanoBRET ratios of the control samples from the experimental ligand-containing samples. We simultaneously fitted the total and nonspecific saturation binding curves using the equation following manufacturer’s instructions.

### High-throughput RNA sequencing and bioinformatics

Stranded libraries of cDNA were prepared by random priming followed by either enrichment in poly(dA) with an oligo(dT) resin or depletion from ribosomal RNA. A minimum of 30 Gb sequencing was obtained for each sample with reads of 150 bases. All poly(dA)-enriched libraries were sequenced on a same flow cell. Reads were aligned with STAR 2.5.0a (Dobin et al. 2013) on human genome version Hg19 primary assembly without patches, allowing only single alignments. Transcriptome analysis on poly(dA)-enriched libraries was performed with edgeR 3.16.5 (Robinson et al. 2009) and raw counts were corrected with a TMM normalization. Divergent transcription sites were extracted from the previous alignments by identifying region in between groups of reads mapping on opposite strands separated by less than 500 bp, using python scripts and bedtools 2.27.1. Reads inside HERV sequences and chromatin states were quantified with featureCount 1.6.0 from the Subread package (Liao et al. 2014). Identification of genes in the neighborhood of HERVs was carried out with GREAT (McLean et al. 2010). Gene ontology analysis was carried out with ENRICHR (Kuleshov et al. 2016).

## Acknowledgements

We thank S El Messaoudi-Aubert for valuable discussion and Jacob Seeler for critical reading of the manuscript. The work supported by grants from REVIVE Agence National de la Recherche and Global care.

---

**Figure S1:** Dieldrin is an activator of several MS-HERVs

(A) Description and references linking HERVs to MS. (B) To estimate the number of loci amplified by the primers we, digital droplet PCR was carried out on Jurkat cell DNA, using DNA from a haploid cell line as a reference. (C-D) Jurkat T cells were treated with either DMSO (vehicle) or with 100µM of bpV(pic) for the indicated times. Abundance of mRNA for the indicated genes or HERVs was assessed by RT-qPCR. Data shown are means +/-SEM from three independent experiments. Significance (p-value) was estimated using the two-sided student t test. (E) Impact of dieldrin on cell viability at the indicated times. (F-G) Jurkat T cells were treated with either DMSO (vehicle) or with 100µM of either Dieldrin or PMA for the indicated times. Abundance of mRNA for the indicated genes was assessed by RT-qPCR. Data shown are means +/-SEM from three independent experiments. Significance (p-value) was estimated using the two-sided student t test. (H) NIH3T3 cells were treated with either DMSO (vehicle), 100µM dieldrin, or 0.1µM topoisomerase inhibitor camptothecin for the 60 min. Cells were fixed, permeabilized, and used for indirect immunofluorescent staining with the indicated antibodies.

**Figure S2:** Dieldrin activates multiple signal transduction pathways.

(A) A human phospho-kinase antibody array was probed with extracts from Jurkat cells treated with either DMSO, dieldrin, or PMA for 30 min. as indicated. The array was processed as a western and revealed by ECL. Spots are distributed as indicated by the provider of the array (https://resources.rndsystems.com/pdfs/datasheets/ary003b.pdf). (B) Spots in the array shown in A were quantified using Fiji (Schindelin et al. 2012). (C-G) Jurkat cells were treated for 30 min. with either DMSO (vehicle) or 100µM Dieldrin in the absence or in the presence of the MEK inhibitor PD98059. Abundance of mRNA for the indicated genes or HERVs was assessed by RT-qPCR. Data shown are means +/-SEM from three independent experiments. Significance (p-value) was estimated using the two-sided student t test.

**Figure S3:** Dieldrin interferes with H3K9me3 and HP1-mediated transcriptional silencing

(A-E) Jurkat cells were treated for 16 hours with either DMSO (vehicle), the EHMT2/G9a-inhibitor BIX-01294, or the DNMT1-inhibitor azacytidine as indicated. (F) Jurkat cells were treated for indicated times with either DMSO (vehicle) or 100µM Dieldrin. In all panels, abundance of mRNA for the indicated genes or HERVs was assessed by RT-qPCR. Data shown are means +/-SEM from three independent experiments. Significance (p-value) was estimated using the two-sided student t test.

**Figure S4:** Transcriptome analysis of Jurkat cells exposed to dieldrin for 30 min.

(A-B) Validation on selected genes activated more by PMA than by dieldrin: Jurkat cells were treated for indicated times with either DMSO (vehicle) or 100µM of either dieldrin or PMA. Abundance of mRNA for the indicated genes or HERVs was assessed by RT-qPCR. Data shown are means +/-SEM from three independent experiments. Significance (p-value) was estimated using the two-sided student t test.

**Figure S5:** HERV transcripts originate from enhancers and from transcription of gene neighborhoods.

(A) Example illustrating the difference between annotation of HERVs by RepeatMasker and HERVd. Location: chr2:52.492.000 on Hg19. (B-D) Screenshots from IGV. Arrows indicate sites of divergent transcription activated by dieldrin. Transcription factor peaks are from the ENCODE Regulation Track of the UCSC Genome Browser. (E) GO term analysis with ENRICHR of genes in proximity (GREAT default settings) of sites of divergent transcription upregulated at least 2-fold in the presence of dieldrin and overlapping with a HERV annotated in HERVd. (F) Example of a transcription factor in a HERV-free zone. Note also that elongation of promoter RNAs (red) and of the main transcript (blue) is increased by dieldrin. (G) Most transcription factor genes have a low density in HERVs: Density of HERVs was evaluated inside a region spanning from −11Kb of the TSS to +11Kb of the TTS. Genes were then segregated in transcription factor (tf) and non-transcription factor genes (not_tf). Graphs indicates the % of genes from each of the two categories fitting into the indicated density bins.

