## Supplementary Materials for "Compound signaling activates endogenous retroviruses by inducing enhancer and gene-neighborhood transcription"

**A**

| Family | GenBank Ref | Ref | Pseudo | repFamily | repClass | repName |
|---|---|---|---|---|---|---|
| HERV_H | AJ289711.1 | Laska and *al*, 2012 | ENV59 | LTR | ERV1 | HERVH-int |
| | AJ289710.2 | | ENV60 | LTR | ERV1 | HERVH-int |
| | AJ289709.1 | | ENV62 | LTR | ERV1 | HERVH-int |
| | | Jern and *al*, 2005 | Pol62 | LTR | ERV1 | HERVH-int |
| HERV_K (HML-2) | AF333072.2 | Tai and *al*, 2008 | K18 | LTR | ERV1 | HERVK-int |
| | AF164610.1 | Christensen, 2005 | K102 | LTR | ERV1 | HERVK-int |
| HERV_W | AF072506.2 | Mameli and *al*, 2007 | ERVWE1 | HERV17-int/LTR17 | ERV1 | HERVW-int |

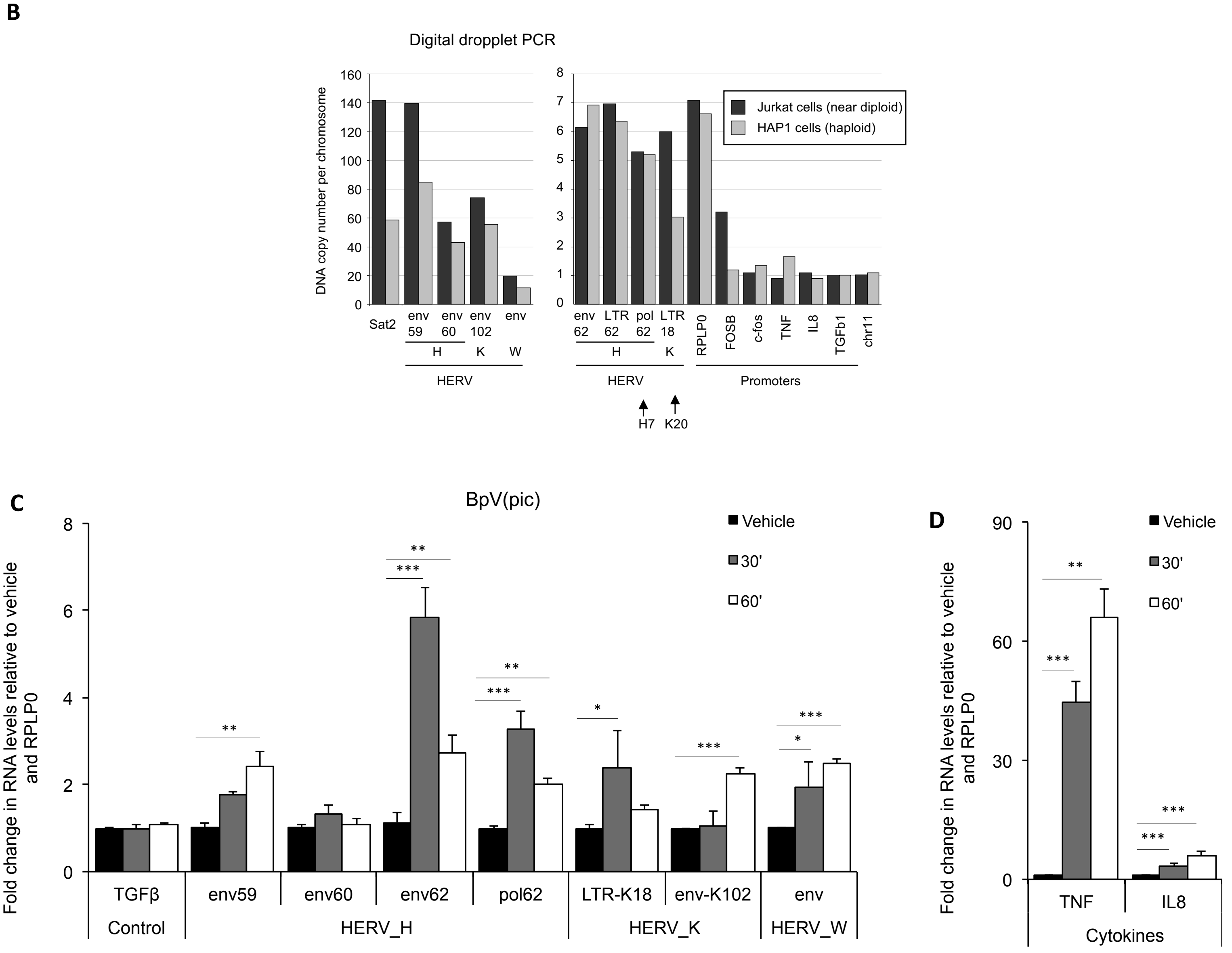


**Figure S1**

**E**

| | Live cells/mL | Dead cells % |
|---|---|---|
| DMSO 1h | 87.104 | 0% |
| Dieldrin in DMSO 30 min. | 85.104 | 0% |
| Dieldrin in DMSO 1h | 92.104 | 2% |

**F**

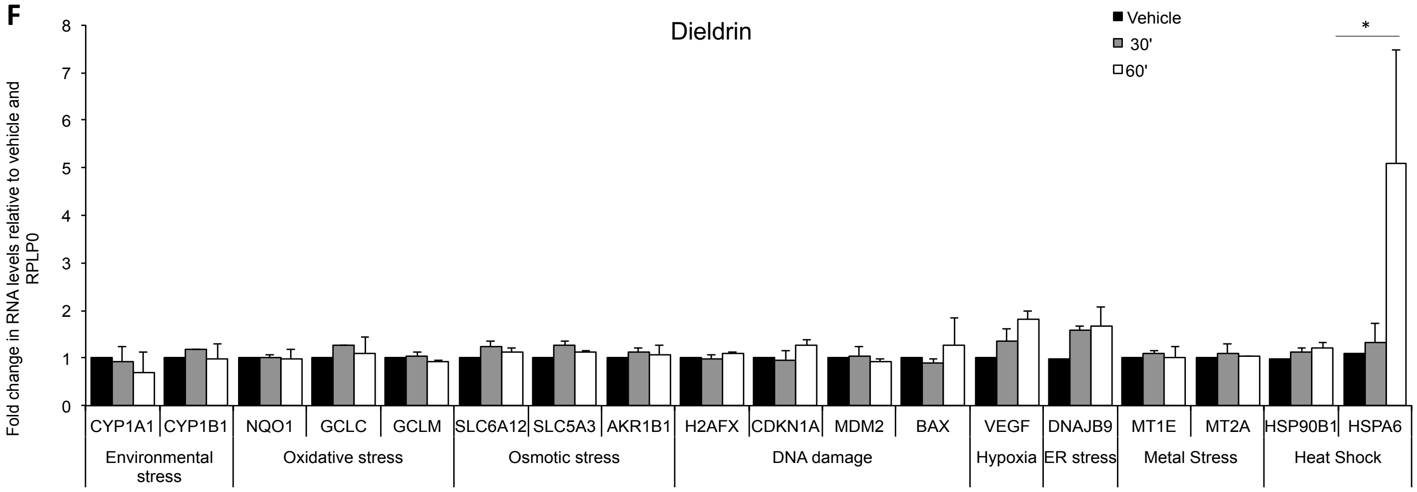


**G**

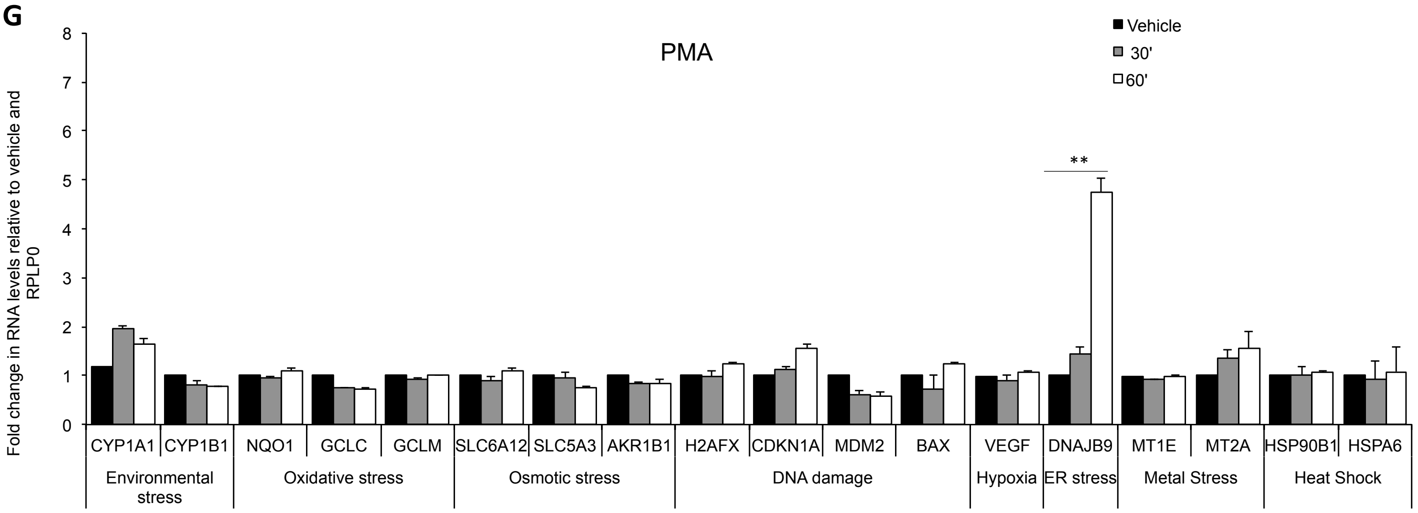


**Figure S1**

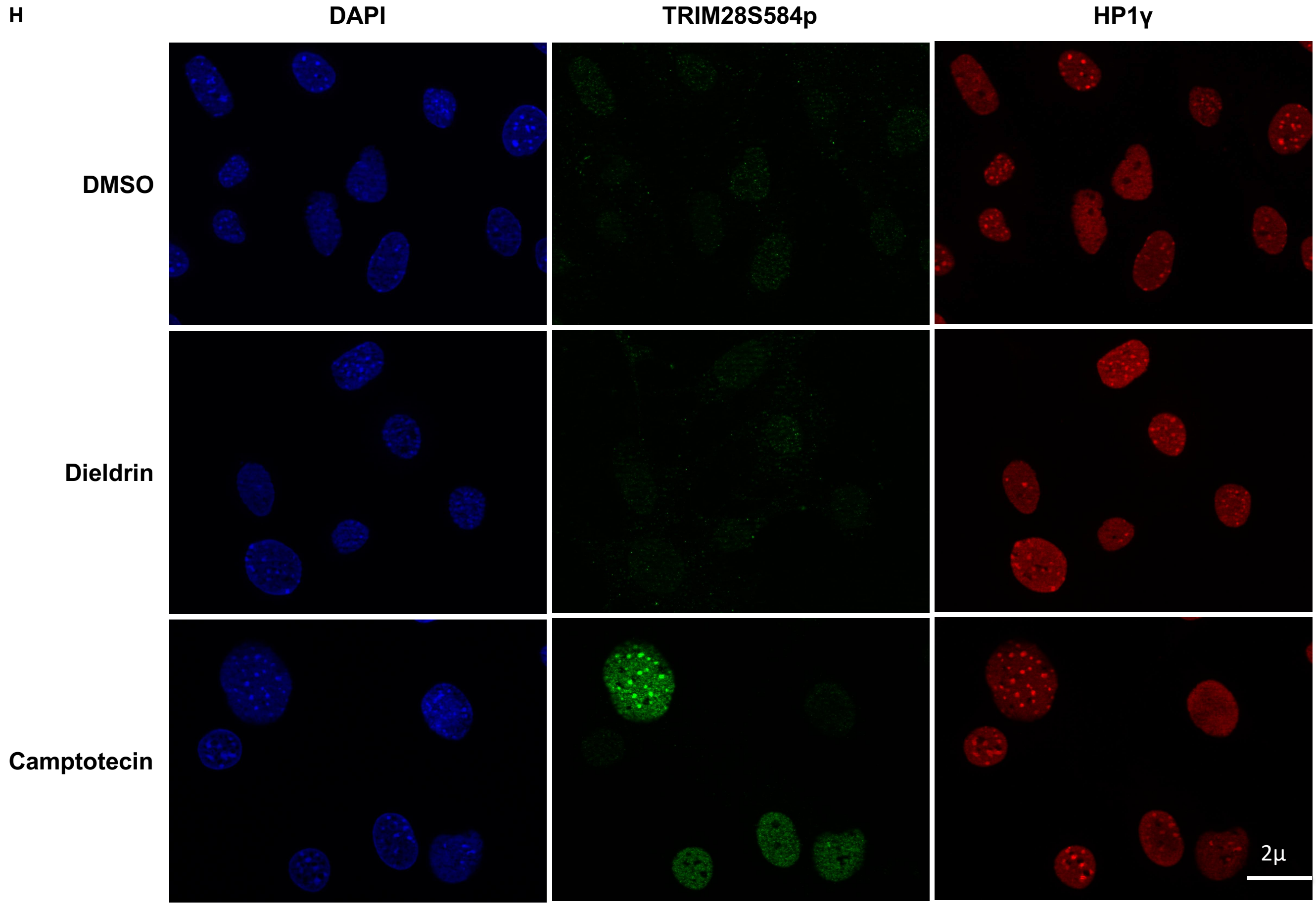


**Figure S1**

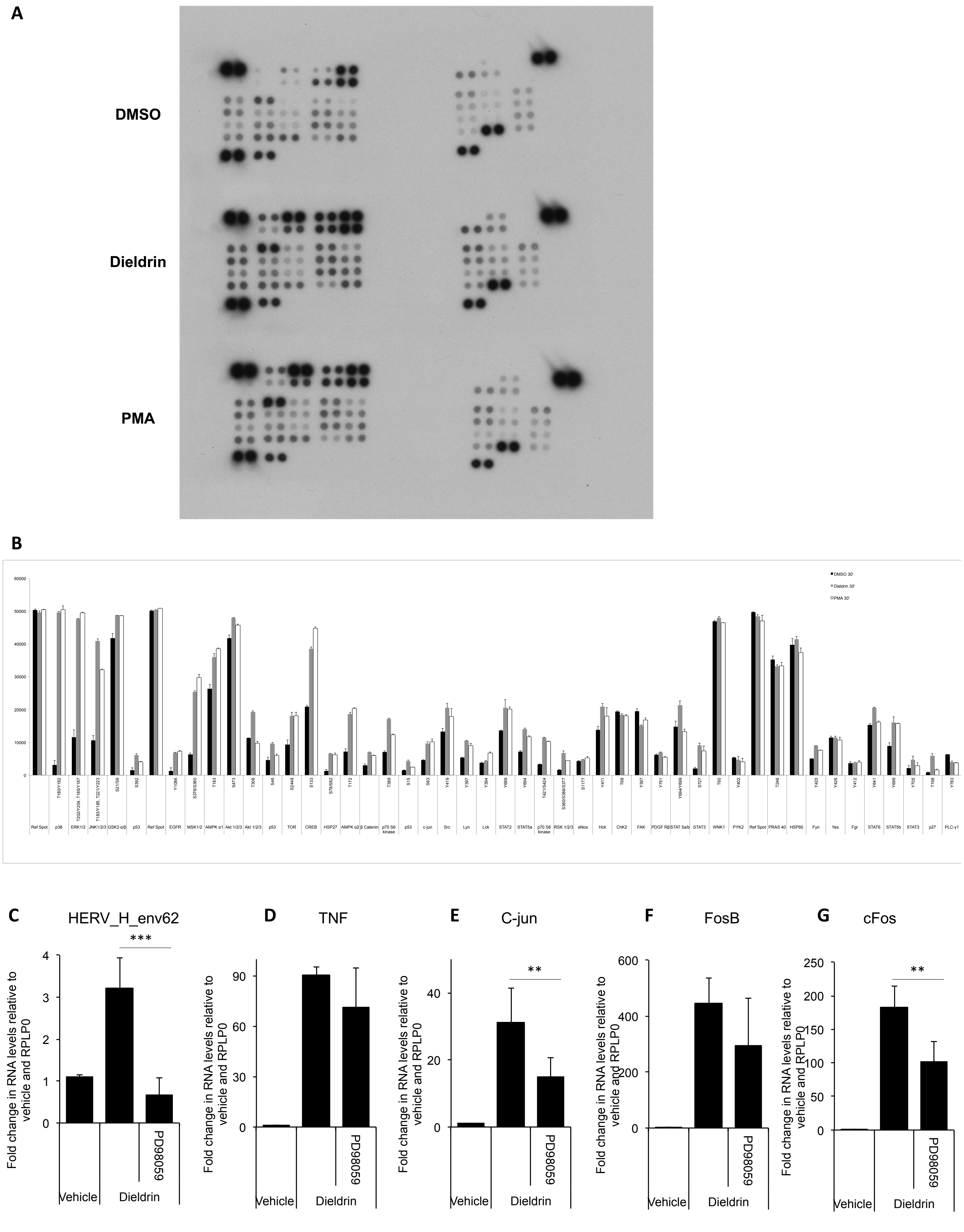

A
DMSO
Dieldrin
PMA
B
C
HERV_H_env62
D
TNF
E
C-jun
F
FosB
G
cFos
Fold change in RNA levels relative to vehicle and RPLP0
Vehicle
Dieldrin
PD98059
***
**
**


**Figure S2**

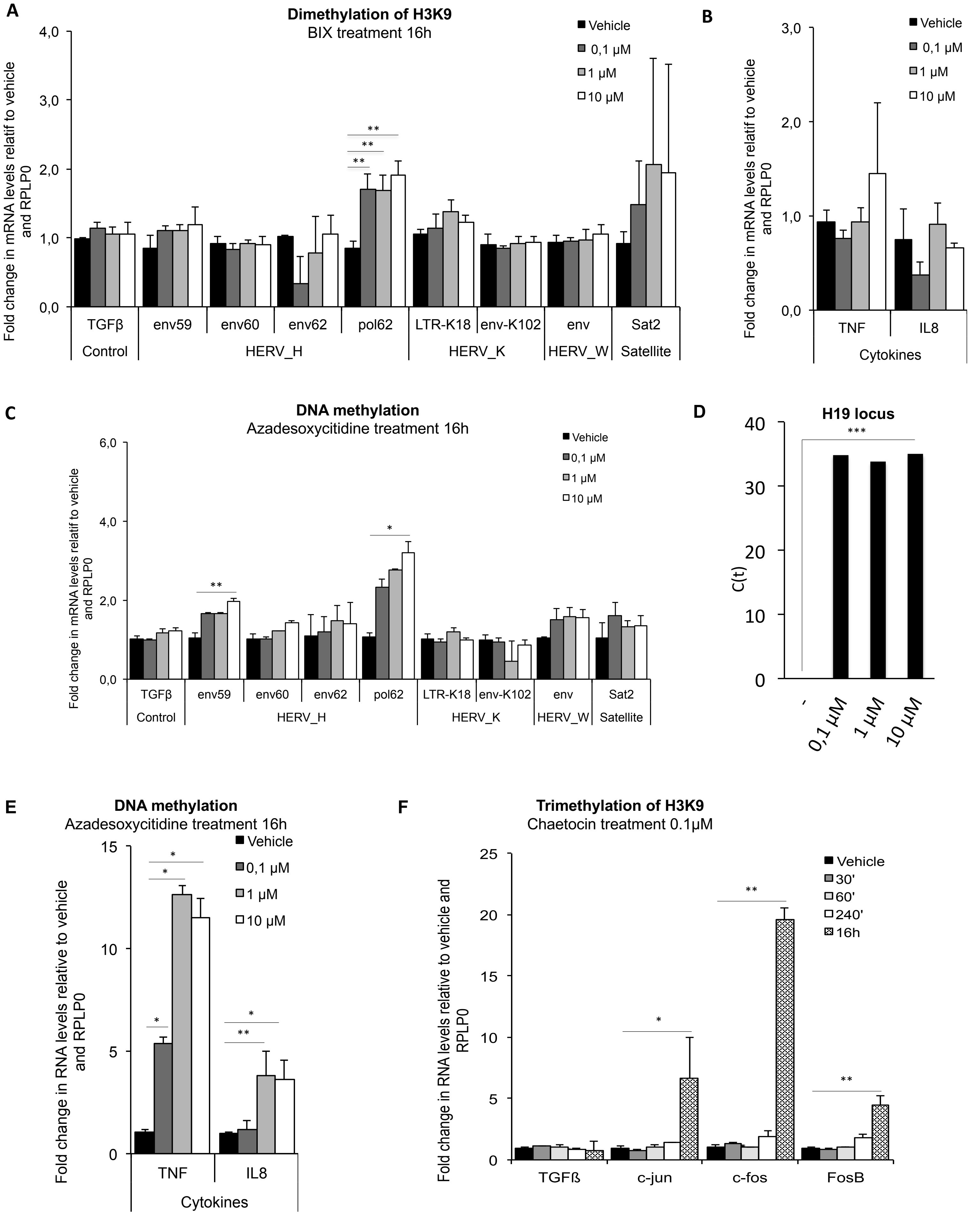

A
Dimethylation of H3K9
BIX treatment 16h
Vehicle
0,1 µM
1 µM
10 µM
Fold change in mRNA levels relatif to vehicle and RPLP0
4,0
3,0
2,0
1,0
0,0
**
**
**
TGFβ
env59
env60
env62
pol62
LTR-K18
env-K102
env
Sat2
Control
HERV_H
HERV_K
HERV_W
Satellite
B
Vehicle
0,1 µM
1 µM
10 µM
Fold change in mRNA levels relatif to vehicle and RPLP0
3,0
2,0
1,0
0,0
TNF
IL8
Cytokines
C
DNA methylation
Azadesoxycitidine treatment 16h
Vehicle
0,1 µM
1 µM
10 µM
Fold change in mRNA levels relatif to vehicle and RPLP0
6,0
4,0
2,0
0,0
**
*
TGFβ
env59
env60
env62
pol62
LTR-K18
env-K102
env
Sat2
Control
HERV_H
HERV_K
HERV_W
Satellite
D
H19 locus
***
C(t)
40
30
20
10
0
-
0,1 µM
1 µM
10 µM
E
DNA methylation
Azadesoxycitidine treatment 16h
Vehicle
0,1 µM
1 µM
10 µM
Fold change in RNA levels relative to vehicle and RPLP0
15
10
5
0
*
*
*
*
**
TNF
IL8
Cytokines
F
Trimethylation of H3K9
Chaetocin treatment 0.1µM
Vehicle
30'
60'
240'
16h
Fold change in RNA levels relative to vehicle and RPLP0
25
20
15
10
5
0
*
**
**
TGFß
c-jun
c-fos
FosB


**Figure S3**

A

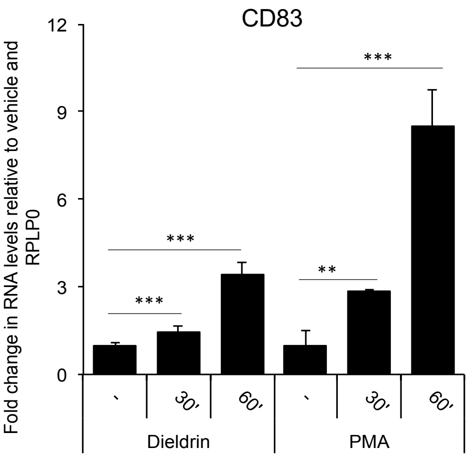


B

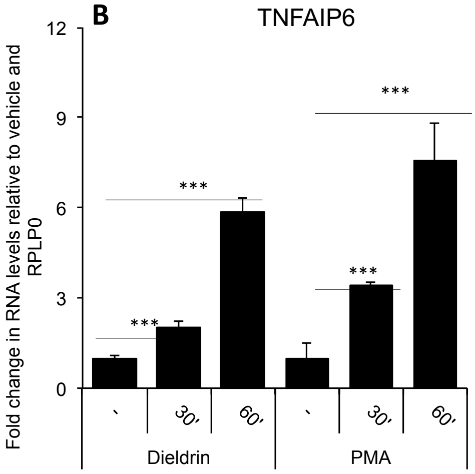


C

Table showing values for repressed genes mentioned in the text

D

Panel showing that when reads with 20As overlap with HERVs, their A tract is mostly encoded by the geneome.

**Figure S4**

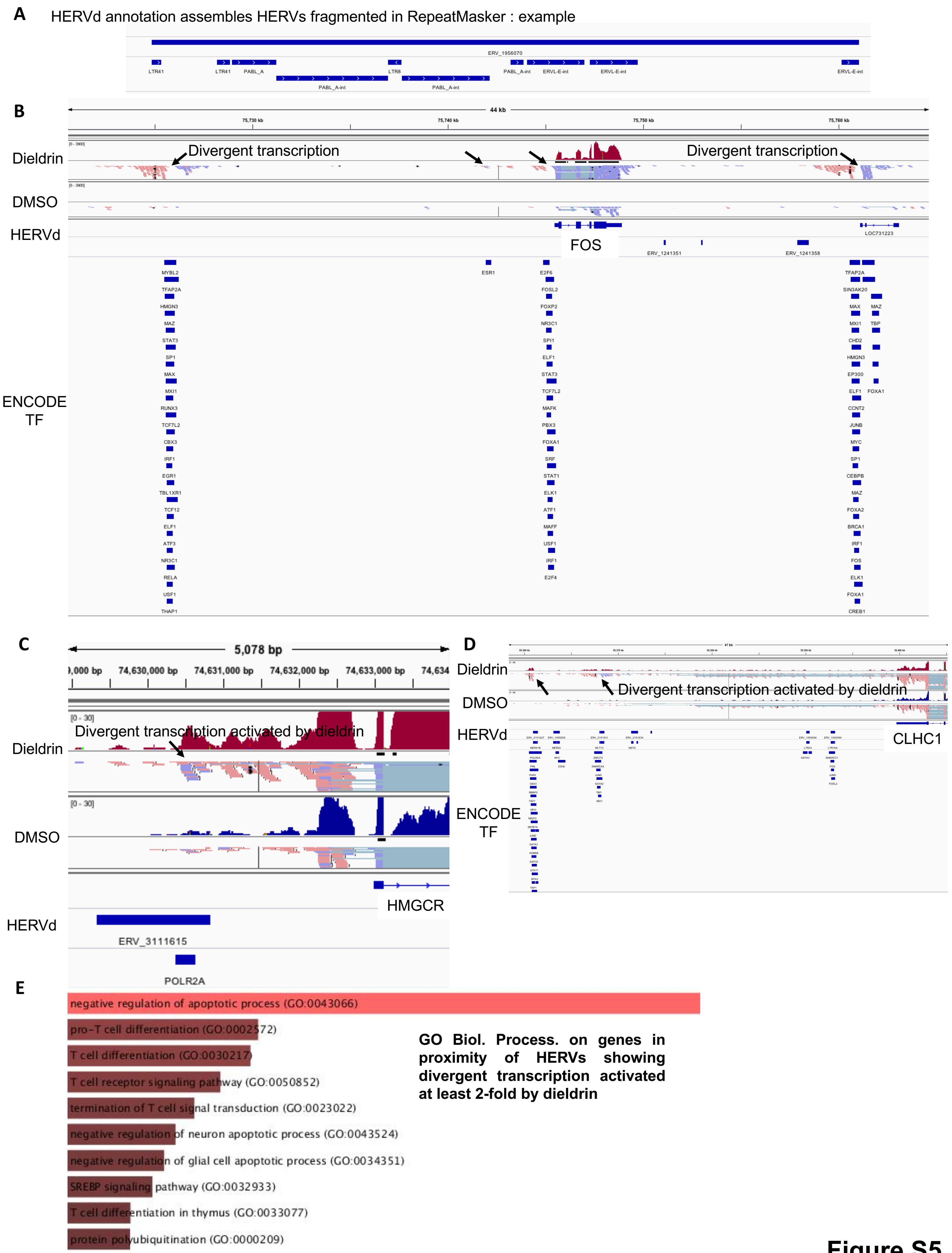
A
HERVd annotation assembles HERVs fragmented in RepeatMasker : example
ERV_1956070
LTR41
LTR41
PABL_A
LTR8
PABL_A-int
ERVL-E-int
ERVL-E-int
ERVL-E-int
PABL_A-int
PABL_A-int
B
44 kb
Dieldrin
Divergent transcription
Divergent transcription
DMSO
HERVd
FOS
LOC731223
ERV_1241351
ERV_1241358
ENCODE TF
MYBL2
TFAP2A
HMGN3
MAZ
STAT3
SP1
MAX
MXI1
RUNX3
TCF7L2
CBX3
IRF1
EGR1
TBL1XR1
TCF12
ELF1
ATF3
NR3C1
RELA
USF1
THAP1
ESR1
E2F6
FOSL2
FOXP2
NR3C1
SPI1
ELF1
STAT3
TCF7L2
MAFK
PBX3
FOXA1
SRF
STAT1
ELK1
ATF1
MAFF
USF1
IRF1
E2F4
TFAP2A
SIN3AK20
MAX
MAZ
MXI1
TBP
CHD2
HMGN3
EP300
ELF1
FOXA1
CCNT2
JUNB
MYC
SP1
CEBPB
MAZ
FOXA2
BRCA1
IRF1
FOS
ELK1
FOXA1
CREB1
C
5,078 bp
74,630,000 bp
74,631,000 bp
74,632,000 bp
74,633,000 bp
Dieldrin
[0 - 30]
Divergent transcription activated by dieldrin
DMSO
[0 - 30]
HMGCR
HERVd
ERV_3111615
POLR2A
D
Dieldrin
DMSO
Divergent transcription activated by dieldrin
HERVd
CLHC1
ENCODE TF
E
negative regulation of apoptotic process (GO:0043066)
pro-T cell differentiation (GO:0002572)
T cell differentiation (GO:0030217)
T cell receptor signaling pathway (GO:0050852)
termination of T cell signal transduction (GO:0023022)
negative regulation of neuron apoptotic process (GO:0043524)
negative regulation of glial cell apoptotic process (GO:0034351)
SREBP signaling pathway (GO:0032933)
T cell differentiation in thymus (GO:0033077)
protein polyubiquitination (GO:0000209)
GO Biol. Process. on genes in proximity of HERVs showing divergent transcription activated at least 2-fold by dieldrin

**Figure S5**

**F** Example of a transcription factor gene in HERV-free zone: **MYC**. Note the increased elongation in the presence of dieldrin.

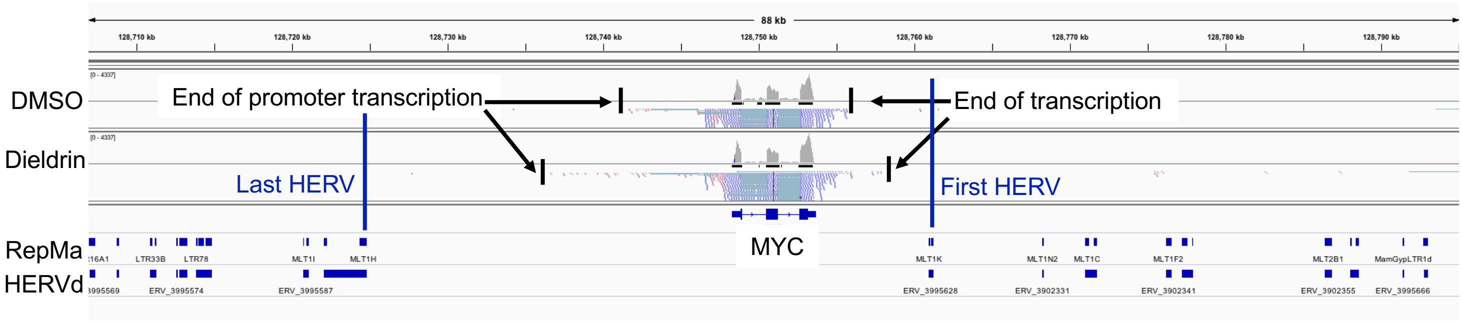


**G** Distribution of HERV densities in transcription factor genes and other genes (in %)

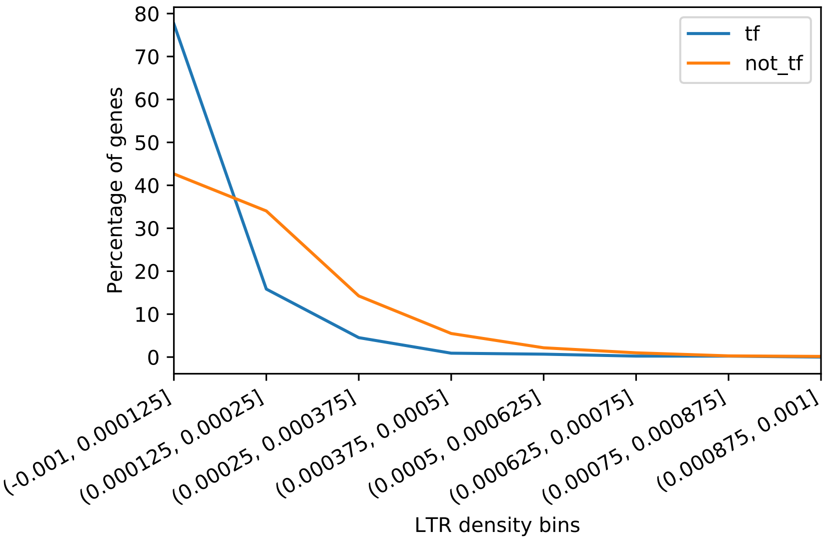


**Figure S5**
